# The CBP KIX domain regulates long-term memory and circadian activity

**DOI:** 10.1101/2020.06.08.130815

**Authors:** Snehajyoti Chatterjee, Christopher C. Angelakos, Ethan Bahl, Joshua D. Hawk, Marie E. Gaine, Shane G. Poplawski, Anne Schneider-Anthony, Manish Yadav, Giulia S. Porcari, Jean-Christophe Cassel, K. Peter Giese, Jacob J. Michaelson, Lisa C. Lyons, Anne-Laurence Boutillier, Ted Abel

## Abstract

CREB-dependent transcription necessary for long-term memory is driven by interactions with CREB-binding protein (CBP), a multi-domain protein that binds numerous transcription factors. Identifying specific domain functions for multi-action proteins is essential to understand processes necessary for healthy living including cognitive function and a robust circadian clock. We investigated the function of the CBP KIX domain in hippocampal memory and gene expression using CBP^KIX/KIX^ mice with mutations that prevent phospho-CREB (Ser133) binding. We found that CBP^KIX/KIX^ mice were impaired in long-term, but not short-term spatial memory in the Morris water maze. Using an unbiased analysis of gene expression after training for hippocampus-dependent memory, we discovered dysregulation of CREB and CLOCK target genes and downregulation of circadian genes in CBP^KIX/KIX^ mice. With our finding that the CBP KIX domain was important for transcription of circadian genes, we profiled circadian activity in CBP^KIX/KIX^ mice. CBP^KIX/KIX^ mice exhibited delayed activity peaks after light offset and longer free-running periods in constant dark, although phase resetting to light was comparable to wildtype. These studies provide insight into the significance of the CBP KIX domain by defining targets of CBP transcriptional co-activation in memory and the role of the CBP KIX domain *in vivo* on circadian rhythms.

## Introduction

Phosphorylation and activation of the cyclic-AMP response element-binding protein (CREB) is fundamental for the induction of gene transcription during long-term memory consolidation (1–3). Decreased CREB levels impair spatial memory whereas overexpression of CREB in the dorsal hippocampus enhances memory (1, 4, 5). However, CREB-induced transcription during memory formation appears tightly regulated as mice with constitutively active hippocampal CREB exhibit impairments in the retrieval of spatial memory (6). Dysregulation of cAMP-PKA signaling and alterations in CREB activity have been associated with age-related cognitive impairments and neurodegenerative diseases (7). In recent U.S. National Health Surveys, more than 46% of respondents over age 65 reported memory impairments (8). Currently many countries around the world, including the United States, are experiencing demographic shifts towards older populations. By 2050, adults 65 and older are predicted to comprise 16% of the world’s population, more than 1.5 billion individuals (9). For neurodegenerative diseases such as Alzheimer’s disease, individuals frequently experience mild cognitive impairments and memory issues years prior to disease diagnosis. Given that increased longevity increases the individual risk and societal economic burden of age-related diseases (10), there is a crucial need to understand the mechanisms and processes involved in long-term memory consolidation.

During memory consolidation, new gene expression is temporally regulated following learning, resulting in specific patterns and waves of transcription (11). Regulation of CREB activity occurs, at least in part, in response to cAMP signaling and PKA phosphorylation of Ser-133. Following this phosphorylation, the kinase inducible domain (KID) of CREB binds to the KIX domain of the cyclic-AMP response element-binding protein (CREB) binding protein (CBP) through an induced fit mechanism (12, 13). The interaction of phosphorylated CREB with the CBP KIX domain appears to be a specific response to cAMP-PKA signaling, and it can be modulated by external stimuli resulting in transcriptional specificity in the expression of target genes (12). As CREB phosphorylation may also occur in response to kinases independent of cAMP-PKA signaling (14), one method through which downstream target genes in memory may be identified is through the manipulation of CREB-CBP interactions at the KIX domain. CBP is a large 265 kDa protein containing multiple interaction domains, including the nuclear hormone receptor binding domain, transitional adapter zinc finger domains, KIX domain, bromodomain, histone acetyltransferase domain, and glutamine-rich (Q) domain. CBP interacts with numerous transcription factors to potentially regulate 16,000 genes (15–18). In hippocampus-dependent memory, *Cbp* mutants exhibit deficiencies in contextual fear conditioning and object recognition memory (19–21). Moreover, CBP functions as a histone acetyltransferase impacting gene expression during memory consolidation (22–27), and more specifically in memory encoding in the medial prefrontal cortex (26, 28). While restoring CREB activity in the CA1 of the dorsal hippocampus enables recovery from spatial memory deficits in an amyloid beta model of Alzheimer’s disease (29), pharmacological activation of CBP/p300 HAT function also improves spatial learning in wildtype (WT) mice and restores spatial long-term memory retention and hippocampal plasticity in a tauopathy mouse model (30, 31). Decreased levels or dysregulation of CBP have also been associated with neurodegenerative diseases including Huntington’s Disease (32, 33) and Alzheimer’s disease (34–36). Previous research on Alzheimer’s disease using rodent models found that decreases in hippocampal CBP activity levels are accompanied by decreased CREB activation, i.e. phospho-CREB levels at Ser-133, although overall CREB levels were not changed (34), emphasizing the need to understand the function of CBP in transcription and memory.

As a far-reaching co-activator of transcription, CBP also regulates the endogenous circadian clock (37). The circadian clock coordinates tissue specific transcriptional regulation of clock-controlled genes through the core circadian transcription factors BMAL1 and CLOCK. CBP is recruited by the CLOCK/BMAL1 complex and putatively interacts with BMAL1 (38, 39). CBP overexpression has been shown to increase CLOCK/BMAL1 mediated transcription of the circadian gene *period1 (per1)* (40). Despite previous identification of CBP interactions with circadian transcription factors, the role of CBP in circadian behavior has not been characterized *in vivo*.

In this study, we used mice (CBP^KIX/KIX^ mice) expressing CBP with three point mutations in the KIX domain of CBP, thereby preventing either phospho-CREB or c-Myb binding to this region (41) to identify the downstream transcriptional pathways and targets in memory regulated by CBP KIX domain interactions, and to characterize the role of CBP KIX domain interactions on circadian activity. CBP^KIX/KIX^ mice were previously shown to present deficits in long-term memory, related to contextual fear conditioning and novel object recognition (19, 42); however, neither the transcriptional profile of these mice after learning nor their circadian behavior has been studied. We found that CBP^KIX/KIX^ mice exhibit specific deficits in long-term spatial memory in the Morris water maze (MWM), although no impairments were observed in these mice for task acquisition or short-term spatial memory. Given the similar memory impairments found for MWM and contextual fear conditioning, we performed deep RNA sequencing from hippocampal tissue following learning for contextual fear conditioning and MWM in CBP^KIX/KIX^ mice and WT littermates to identify downstream gene targets of transcriptional co-activation through the KIX domain. Pathway analysis suggested that CREB was the prominent upstream regulator of the differentially expressed genes between CBP^KIX/KIX^ mice and WT mice. Circadian clock related genes were among the most deregulated genes after learning in the hippocampus of CBP^KIX/KIX^ mice. We characterized circadian rhythms in CBP^KIX/KIX^ mice and found that they have a lengthened free-running circadian period compared to WT littermates. Surprisingly, the ability to phase shift activity in response to light pulses was retained in CBP^KIX/KIX^ mice. These studies provide significant insight into the role of phospho-CREB-CBP interactions in the regulation of learning-induced transcription, memory consolidation, and circadian rhythms.

## Results

### CBP^KIX/KIX^ mice are deficient in long-term but not short-term spatial memory in the Morris water maze

Previously, the KIX domain of CBP, important for protein-protein interactions (13, 43), was found to be essential for long-term memory after contextual fear conditioning (19) and training in the novel object recognition task (42). However, studies have identified additional CBP functions that contribute to long-term memory including histone acetylation (30, 44). To identify the specific role of the KIX domain in hippocampus-dependent long-term memory, we analyzed the consequences of a CBP KIX domain mutation on short and long-term spatial memory using the MWM, a hippocampus-dependent task. Unlike other learning paradigms, MWM requires multiple days of training and is considered the gold standard of spatial memory tasks (45). CBP^KIX/KIX^ mice and their WT littermates were trained over 5 consecutive days to locate a hidden platform positioned at a fixed location. Independent groups of mice were tested at separate time points to assess retention, either 1 h after the last training for short-term memory or 24 h after training for long-term memory (**Fig 1a**). Both WT and CBP^KIX/KIX^ mice showed a day-to-day decrease (D1 to D5) in escape latencies indicating significant learning of the task (**Fig 1b**). CBP^KIX/KIX^ mice demonstrated an overall improvement throughout training similar to their WT littermates. These results suggest that CBP^KIX/KIX^ mice can improve their performance in MWM even without transcription factor interaction through the KIX domain of CBP. We assessed learning retention during the probe test by comparing the time spent in the target quadrant with time spent in the other three quadrants. During the short-term memory test (1 h), (WT, n=10; CBP^KIX/KIX^, n=6, **Fig 1c**), WT control mice showed significant memory of the platform location with more time spent in the target quadrant compared to search time in other quadrants. Similar to WT, CBP^KIX/KIX^ mice explored significantly more in the target quadrant compared to the other three quadrants during the short-term probe test with the search time above the chance time of 15s in the target quadrant (WT: 34.9s and CBP^KIX/KIX^: 29.0s in target quadrant (**Fig 1c**)). However, during the long-term memory probe test (WT, n=10; CBP^KIX/KIX^, n=6, probe test 24h, **Fig 1d**), WT mice exhibited significant long-term memory with increased search time in the target quadrant over the other three quadrants (WT: 25.2s vs chance 15s), while CBP^KIX/KIX^ mice searched randomly in the four quadrants reflecting the lack of long-term memory (CBP^KIX/KIX^: 14.9s vs chance 15s). These results indicate that CBP^KIX/KIX^ mice have deficits in long-term spatial memory consolidation, consistent with previous research on contextual memory (19), and suggest a crucial role for the interaction of the KIX domain with transcription factors during this process.

**Figure 1.**
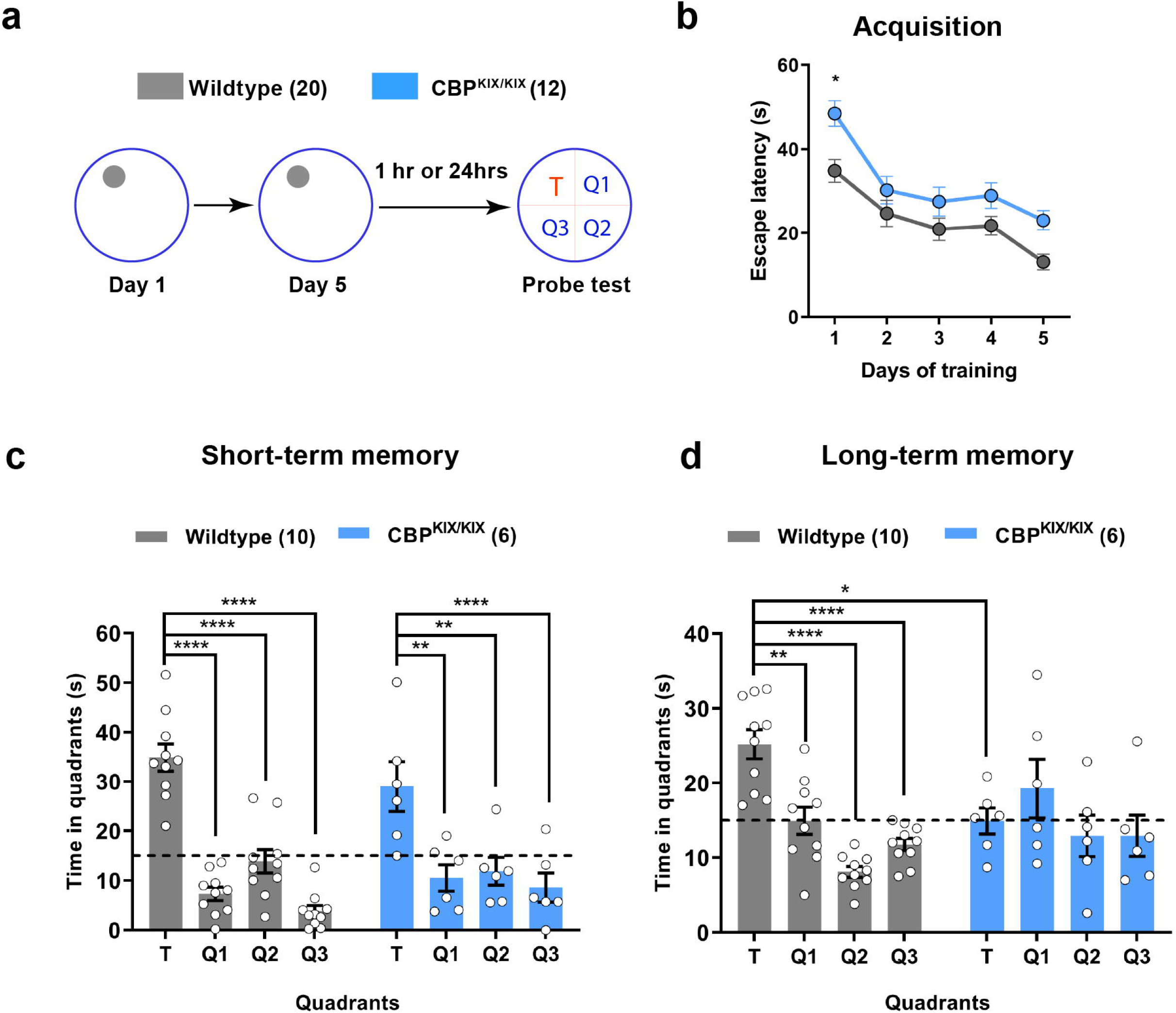
CBP^KIX/KIX^ mice are impaired in long-term spatial memory but have normal short-term retention. **(a)** Experimental scheme. CBP^KIX/KIX^ (n=12;) and WT littermates (n=20, both males and females) were trained in MWM for 5 days. Short-term or long-term memory was assessed either 1 h or 24 h respectively after the last training day. **(b)** The acquisition curve during training indicates that CBP^KIX/KIX^ mice and WT littermates learned the MWM task finding the hidden platform more quickly on subsequent training days [Day effect, F_(3.429, 106.3)_ = 20.57, *p*<0.0001, Day x Genotype interaction, ns] **(c)** In the 1 h probe test for short-term memory, both CBP^KIX/KIX^ (n=10) and WT mice (n=6) showed significantly higher preference for the target quadrant suggesting intact short-term retention. Bar graphs are mean ± SEM. Two-way ANOVA revealed no interaction between “genotype” and “quadrant location” F _(3, 42)_ = 1.286, *p*=0.2917. Significant main effect of quadrant location was observed F _(3, 42)_ = 29.07, *p*<0.0001, while no effect of genotype was seen F _(1, 14)_ = 1.000, *p*=0.3343. Sidak multiple comparisons revealed no significant difference between WT and CBP^KIX/KIX^ mice in the time spent in target quadrant. **(d)** In the 24 h probe test, CBP^KIX/KIX^ mice (n=6) were impaired in long-term memory as shown by reduced exploration in the target quadrant compared to WT littermates (n=10). 2-way ANOVA revealed significant interaction between “genotype” and “quadrant location” F _(3, 42)_ = 4.476, *p*=0.0082 and a main effect of quadrant location F _(3, 42)_ = 6.835, *p*=0.0007. Importantly, Sidak multiple comparisons revealed a significant difference in time in target platform between WT and CBP^KIX/KIX^ mice (adjusted p=0.0030) Data are presented as mean□±□SEM. Differences are significant at \**p*□<□0.05 \*\**p*□<□0.01 \*\*\**p*□<□0.001, evaluated with two-way ANOVA and Sidak multiple correction. TQ, target quadrant; Q1, Q2, Q3 correspond to the three other quadrants.

Hippocampus-dependent learning induces CREB phosphorylation at Ser133, a precursor event for CBP binding. Given that the CBP^KIX/KIX^ mutation prevents phospho-CREB (Ser133) binding to CBP, we hypothesized that phospho-CREB (Ser133) levels would be decreased in CBP^KIX/KIX^ mice as the CREB protein would be more vulnerable to phosphatase activity at the Ser133 site. To test this hypothesis, we performed western blots of protein extracted from WT and CBP^KIX/KIX^ mice 1 h after the third day of MWM training. As expected, we found that MWM increased CREB phosphorylation at Ser133 in WT mice. Phospho-CREB levels were reduced in CBP^KIX/KIX^ mice after training compared to WT mice, although no differences in the baseline levels of CREB phosphorylation at Ser133 were observed in homecage controls (**Supplemental Fig S1**).

### CBP KIX domain mutation alters circadian gene expression in the dorsal hippocampus following spatial learning

As CBP can bind either CREB or c-MYB through the KIX domain (13, 46, 47), we wanted to identify the set of genes for which KIX domain transcription factor binding is necessary to enable hippocampus-dependent memory consolidation. Consequently, we performed an unbiased analysis of gene expression using deep RNA sequencing of dorsal hippocampal tissues from CBP^KIX/KIX^ mice and WT littermates after training in the MWM (RNA-Seq 1) or contextual fear conditioning (RNA-Seq 2) **(Fig 2a).** Spatial memory has been shown dependent upon the dorsal, but not ventral, hippocampus (48–51). As the objective was to identify gene expression common to multiple types of hippocampus-dependent memory consolidation, we combined data from RNA-Seq 1 and RNA-Seq 2. We tested for differences in expression between WT and CBP^KIX/KIX^ mice, using experimental batch (i.e., RNA-Seq 1/RNA-Seq 2) as a covariate to delineate genes dysregulated by the KIX domain across multiple paradigms of hippocampus-dependent learning. We identified 158 differentially expressed genes (DEGs) at a false discovery rate (FDR) of 0.05 (**Fig 2b**).

**Figure 2.**
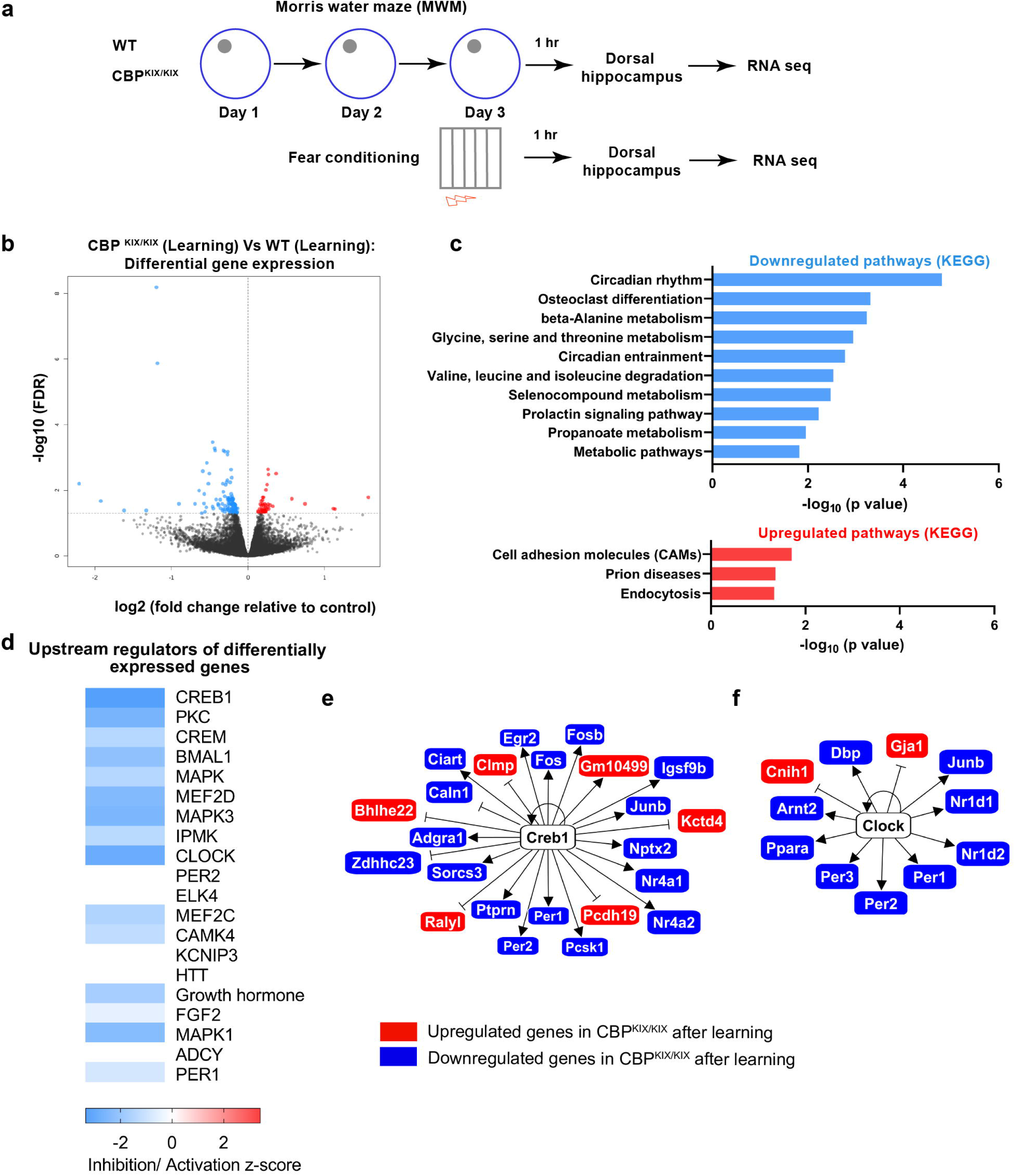
Differential gene expression analysis in the dorsal hippocampus of CBP^KIX/KIX^ mice following spatial learning. **(a)** Experimental scheme: Total RNA was extracted from the dorsal hippocampus of CBP^KIX/KIX^ and control littermates one hour after MWM training (day 3, trial 4) and one hour after contextual fear conditioning (n= 7 CBP^KIX/KIX^, n= 7 controls). Each set of experiments was sequenced separately but analyzed together to identify common genes responsive to hippocampus dependent learning. **(b)** Volcano plot illustrating differentially expressed genes between CBP^KIX/KIX^ and control mice after learning (common between MWM and contextual fear conditioning). **(c)** KEGG pathway enrichment network analysis showing significant KEGG pathways enriched in the downregulated genes. Top significant pathways are shown in the bar graph. Mammalian circadian rhythm pathway (p=1.55×10^−5^) **(d)** Heat map of IPA upstream regulator analysis on differentially expressed genes in CBP^KIX/KIX^ mice following training. Most significant regulators are on the top of the heat map. CREB1 also known as CREB, is the top-predicted upstream regulator of differentially expressed genes (FDR<0.05) in the CBP^KIX/KIX^ mice compared to wildtype littermates (predicted inhibition, z-score= −3.349, p=1.49 × 10^−14^. **(e) and (f),** Known interactions of differentially expressed genes (FDR<0.05) in CBP^KIX/KIX^ vs control following learning predicting CREB1 **(e)** and CLOCK **(f)** as upstream regulators.

Among the DEGs, 114 genes were downregulated while 44 were upregulated in CBP^KIX/KIX^ mice relative to WT **(Supplemental Table 1)**. As CBP positively regulates transcription (52), we hypothesized that the downregulated genes in CBP^KIX/KIX^ mice may have a more direct impact on memory formation. Utilizing enrichment network analysis after annotating the downregulated DEGs through the Kyoto Encyclopedia of Genes and Genomes (KEGG) pathway database, we found the downregulated DEGs to be most significantly enriched for genes in the mammalian circadian rhythm pathway (**Fig 2c**), thus specifically linking the KIX domain of CBP to the regulation of circadian gene transcription and expanding upon previous research establishing interactions between CBP and the circadian transcriptional heterodimer CLOCK-BMAL1 (38, 40, 53, 54). We found that the most significantly upregulated pathway in CBP^KIX/KIX^ mice was the cell adhesion molecules pathway.

To identify the transcriptional drivers of DEGs in CBP^KIX/KIX^ mice in an unbiased fashion, we used QIAGEN’s Ingenuity® Pathway Analysis (IPA). From our observed DEGs, Upstream Regulator Analysis in IPA identified CREB1 as the top predicted upstream regulator of DEGs between CBP^KIX/KIX^ and WT after learning (activation z-score = −3.349) (**Fig 2d, Supplemental Table 2**). Sixteen downregulated and six upregulated genes among the DEGs were found to be regulated by CREB (**Fig 2e**). The second most significant upstream regulator of DEGs was CLOCK (activation z-score = −2.927). BMAL1 (also known as ARNTL), a binding partner of CLOCK, also ranked highly as a significant upstream regulator of DEGs (activation z-score = −2.192). Among the CLOCK regulated genes, we identified genes related to circadian rhythm including *Nr1d1, Nr1d2, Per1, Per2, Per3* and *Dbp* (**Fig 2f**).

### Spatial learning induces CBP downstream genes in WT mice

Following our finding that circadian and activity dependent genes were downregulated after learning in CBP^KIX/KIX^ mice, relative to WT mice, we next analyzed gene expression profiles of hippocampus-dependent learning by comparing the WT learning group from both tasks (contextual fear conditioning and MWM) to the WT homecage controls (Experimental Schematic **Fig 3a**). We identified 135 DEGs at an FDR of 0.05 (**Fig 3b)**. Among the DEGs, 47 genes were downregulated while 88 were upregulated following learning **(Supplemental Table 3)**. To externally validate our results, we tested the list of learning-responsive genes for enrichment of genes previously identified as differentially expressed one hour following *in vivo* chemically induced neuronal activation (55). We found a significant enrichment of these positive controls (P=1.035e-10, odds ratio=3.57; **Supplemental Table 4**), suggesting that transcriptional programs reproducibly regulated by learning in multiple behavioral paradigms share significant overlap with genes regulated by *in vivo* chemical activation of neurons.

**Figure 3.**
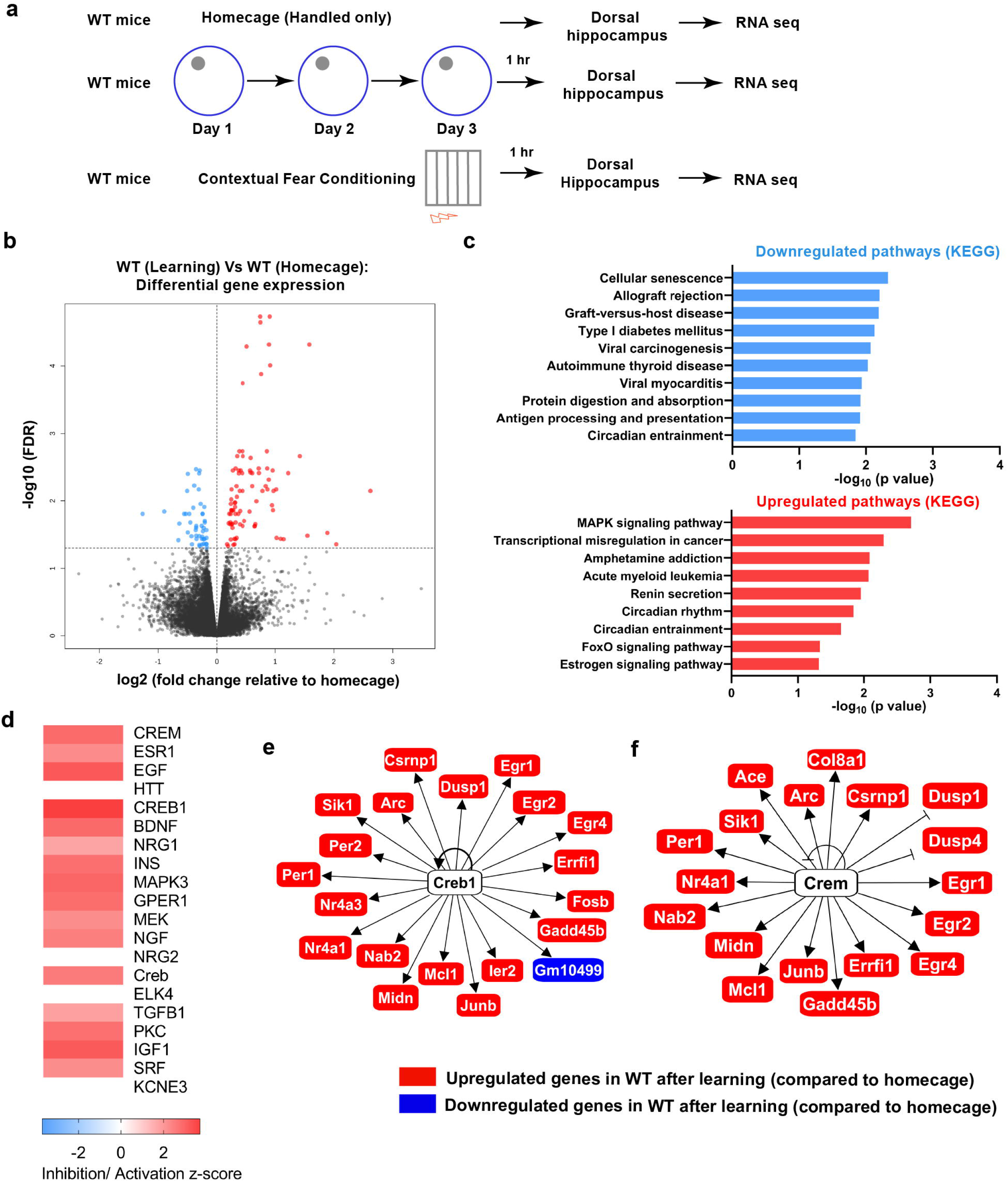
Differential gene expression analysis in the dorsal hippocampus of wildtype mice following learning in fear conditioning and water maze compared to homecage controls. **(a)** Experimental scheme: RNA-Seq analysis performed after learning in Figure 2 was compared to RNA extracted from WT homecage animals. To identify the genes altered by learning, we compared WT contextual fear conditioning (n=4) with WT homecage (n=4) and WT MWM (n=3) with WT homecage (n=3) **(b)** Volcano plot illustrating differentially expressed genes between WT learning and WT homecage mice. **(c)** KEGG pathway enrichment network analysis showing significant KEGG pathways enriched in the upregulated genes. Top significant pathways are shown in the bar graph. MAPK signaling pathway (p= 0.0019) **(d)** Heat map of IPA upstream regulator analysis on differentially expressed genes in CBP^KIX/KIX^ mice following training. Most significant regulators are on the top of the heat map. CREM, is the top-predicted upstream regulator of differentially expressed genes (FDR<0.05) in the WT learning group compared to homecage controls (predicted activation, z-score=2.854, p=2.54 × 10^−21^). CREB1 also appears as an upstream regulator (predicted activation, z-score=3.715, p=3.97 × 10^−13^). **(e)** and **(f)** Known interactions of differentially expressed genes (FDR<0.05) in Learning group vs homecage predicting CREB **(e)** and CREM **(f)** as upstream regulators.

Using enrichment network analysis after annotating the upregulated DEGs through the KEGG pathway database, we found that the most significant change occurred in the MAPK signaling pathway (also known as the Ras-Raf-MEK-ERK pathway) (**Fig 3c**). Circadian genes have recently been shown to be involved in hippocampus-dependent memory (56, 57), and the lack of functional *Per1* in the hippocampus impairs object location memory (58) and LTP (59). As predicted, based upon the downregulation of circadian pathways we observed after learning in CBP^KIX/KIX^ mice, we also found pathways related to circadian rhythm and circadian entrainment to be upregulated following spatial learning in WT mice. Analysis of the most significantly downregulated pathways revealed that genes associated with cellular senescence were the most significantly downregulated genes after learning, consistent with the hypothesis that learning increases neuronal survival with neurogenesis (60–62).

Upstream regulator analysis revealed cAMP-PKA responsive transcription factor CREM and CREB to be the top upstream regulators of the DEGs predicted to be activated following learning (**Fig 3d, Supplemental Table 5**). CREM can be activated through phosphorylation at Ser 117 as well as independently activated in some tissues (63). CREB target genes included 19 upregulated and one downregulated gene (**Fig 3e**), while CREM regulated genes included 18 upregulated genes following learning in WT mice (**Fig 3F**). Interestingly, we found several genes regulated by both CREB and CREM to be upregulated following learning in WT mice and otherwise downregulated in CBP^KIX/KIX^ mice (**Fig 3e and f**; **Fig 2e and f**).

### Comparisons of differential gene expression in hippocampus dependent learning paradigms reflect genes dysregulated after learning in CBP^KIX/KIX^ mice

To integrate the results of the preceding RNA-Seq experiments and to determine whether the learning-responsive genes (MWM and contextual fear conditioning) in WT mice were dysregulated after learning in CBP^KIX/KIX^ mice, we tested whether the genes differentially expressed in CBP^KIX/KIX^ mice following learning were significantly enriched for genes differentially expressed following learning in WT animals compared to the homecage condition. We found a significant enrichment (P=5.91e-10, odds ratio=13.51), suggesting that inhibition of protein interactions through the CBP KIX binding following learning prevents hippocampal learning-regulated transcription. Heat map representations were made based on the twelve most significant DEGs common across data sets **(Fig 4)**. Consistent with our previous analysis, circadian genes were among the most significant genes differentially expressed across data sets with *Per1* and *Per2* in the top upregulated genes in WT mice with learning and downregulated in CBP^KIX/KIX^ mice. Not surprisingly, activity-dependent genes including *Junb, Fosb* and *Nr4a1* were significantly upregulated after learning in WT mice and significantly downregulated in CBP^KIX/KIX^ mice. Sv2b, a synaptic vesicle protein important in vesicle secretion and neurotransmission, was also significantly upregulated in WT mice after learning compared to homecage animals and downregulated in CBP^KIX/KIX^ mice after learning compared to WT mice. Analysis across data sets identified *Pcp4* (Purkinje cell protein 4), a calmodulin binding protein that acts as a modulator of calcium signalling, as significantly downregulated in WT mice after learning and upregulated in CBP^KIX/KIX^ mice after learning suggesting dysregulated intracellular signalling in CBP^KIX/KIX^ mice after learning.

**Figure 4.**
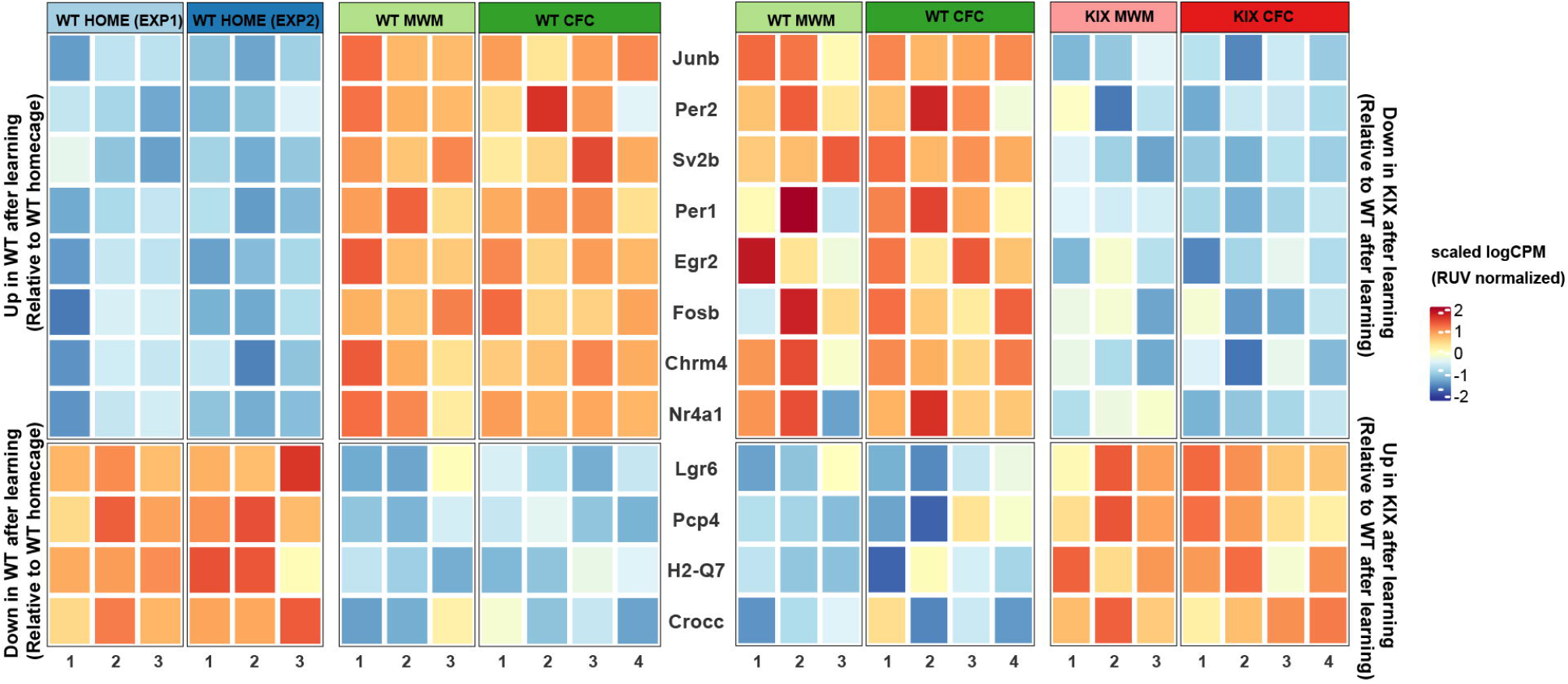
CBP^KIX/KIX^ mice show transcriptional dysregulation of learning-responsive genes. Heat map depicting color-coded expression of the top 12 upregulated or down regulated genes responsive to learning (left) which fail to be appropriately regulated following learning in CBP^KIX/KIX^ mice (right). **Left:** Genes depicted show differential expression between homecage wildtype mice with wildtype mice after learning in the Morris Water Maze (MWM) and contextual fear conditioning (CFC) training. **Right:** The same genes exhibit differential expression between trained wild type mice and trained CBP^KIX/KIX^ mice. Expression values depicted represent the scaled log of RUV-normalized counts per million. As RUV normalization differed between differential expression analyses, scaling was applied independently for each analysis.

### The CBP KIX mutation has only a modest effect on baseline gene expression

In the CBP^KIX/KIX^ mutant mice, it is possible that developmental consequences of the mutation affect gene expression in the hippocampus in such a manner that neural circuitry or basal synaptic transmission would be affected. Although this possibility appears less likely given previous research with these mice in contextual fear conditioning (19) and our behavioral data in the MWM task in which CBP^KIX/KIX^ mice demonstrated task acquisition and short-term memory retention similar to WT mice, we analyzed baseline gene expression in WT and CBP^KIX/KIX^ mice. We performed microarray gene expression analysis using Affymetrix MOE 430v2 arrays in WT and CBP^KIX/KIX^ mice, using RNA extracted from whole hippocampus. We found comparatively few changes in gene expression in CBP^KIX/KIX^ mutant mice in baseline/homecage conditions with almost all of the observed downregulation of gene expression consistent with the function of CBP as a positive regulator of gene expression (**Table 1**). Among the 27 downregulated genes and one upregulated gene that were differentially regulated in the homecage condition between WT and CBP^KIX/KIX^ mice (95% confidence level), *Fos* was the only activity dependent gene downregulated in CBP^KIX/KIX^ mice. Even when the confidence level was reduced to 80%, only 63 genes were found to be downregulated in the mutant mice and three upregulated (**Supplemental Table 6**). Overall, most of the observed gene changes were modest reductions in gene expression (average 23% reduction). The observation of differential expression in such a limited number of genes under homecage conditions is likely a result of the specificity of the KIX mutation as it affects only a single interaction interface of the protein (47). Although unlikely, it is possible that the KIX mutation affected stability of the mRNA encoding CBP. Using three non-overlapping probes, our microarray data confirmed that overall RNA levels for CBP were not reduced in the mutant mice compared to WT mice. The KIX mutation was validated using a fourth CBP probe set that hybridizes with the region containing the mutated base pairs in the CBP^KIX/KIX^ mice. We found that the signal was substantially reduced as predicted (**Supplemental Fig S2**).

**Table 1.**
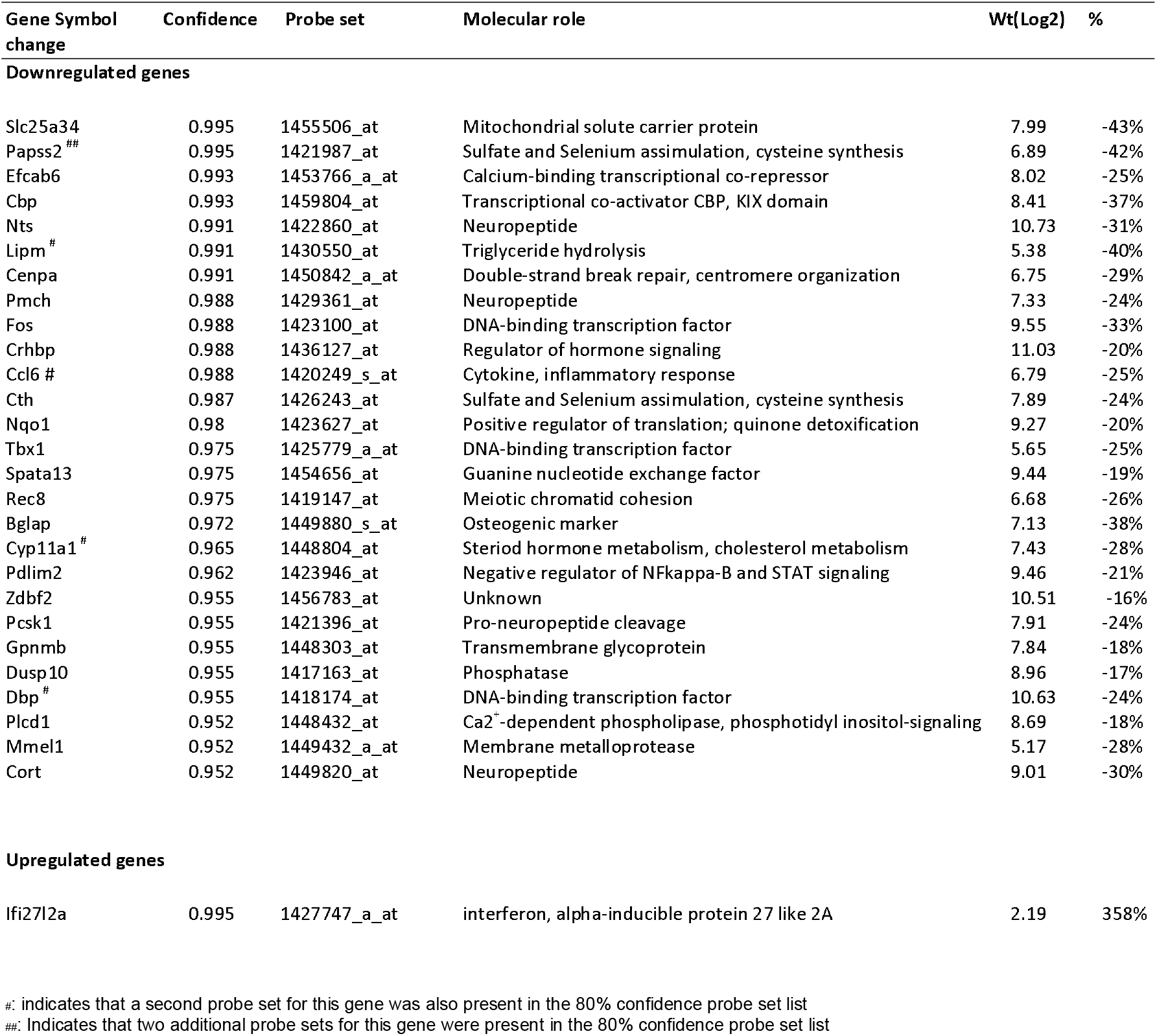
Highest confidence (95%) changes in homecage gene expression in CBP^KIX/KIX^ mice.

### CBP^KIX/KIX^ mice display increased free-running period length and phase differences in peak activity compared to WT mice

Our studies indicate that the CBP KIX domain was important for regulation of gene transcription by the circadian transcription factors CLOCK and BMAL1 even though *in vitro* studies had suggested that CBP interactions with BMAL1 occurred in a region outside of the KIX domain in CBP (64). We then investigated *in vivo* circadian rhythms in CBP^KIX/KIX^ mice. We measured homecage activity in CBP^KIX/KIX^ and WT mice using infrared beam breaks with one week of continuous monitoring in a 12 hour light /12 hour dark (12h:12h LD) cycle. Layered infrared beams allow quantification of locomotor activity as well as rearing, as previously described (65) (**Fig 5a**). CBP^KIX/KIX^ mice had significantly lower anticipatory activity in the last hour of the light (inactive) phase and lower activity in the first two hours of the dark phase. We found that CBP^KIX/KIX^ mice reached peak activity levels three hours later than their WT littermates. Despite the circadian alterations in activity profiles, there was no difference between CBP^KIX/KIX^ and WT mice in total activity across the 24-hour day (**Fig 5b**). We also found no differences in activity when the light and dark phases were analyzed separately (**Fig 5c**). We observed significant delays in activity onset and peak activity in both male and female CBP^KIX/KIX^ mice (**Supplemental Fig 3**). These results indicate that male and female CBP^KIX/KIX^ mice have delayed activity patterns in 12h:12h LD while maintaining normal total locomotor activity levels. Given the temporally shifted activity patterns of CBP^KIX/KIX^ mice, we next investigated whether the CBP^KIX/KIX^ mutation affected free-running circadian rhythms under constant conditions. After entrainment to LD cycles, CBP^KIX/KIX^ and WT mice were placed in constant darkness for 4 weeks to measure circadian activity and period length. CBP^KIX/KIX^ mice had a significantly longer circadian period, ~22.9 minutes, than WT littermates (WT: 23.66 ± 0.01 hrs, CBP^KIX/KIX^ 24.05 ± 0.03 hrs) (**Fig 5d-f**). There was no effect of sex on circadian period with both male and female CBP^KIX/KIX^ mice displaying significantly lengthened free-running periods. This shift in circadian period is roughly similar to the shifts observed in knockouts of core circadian clock genes, including *CLOCK* (66), *Per1* (67), *Per3* (68), *Nr1d1 (69)*, *Dbp* (70), *Rora* (71), *Rorb* (72), and *Npas2 (73)* [summarized in (74)]. Thus, these results provide *in vivo* evidence that the KIX domain of CBP is an integral modulator of circadian period length and activity profiles.

**Figure 5.**
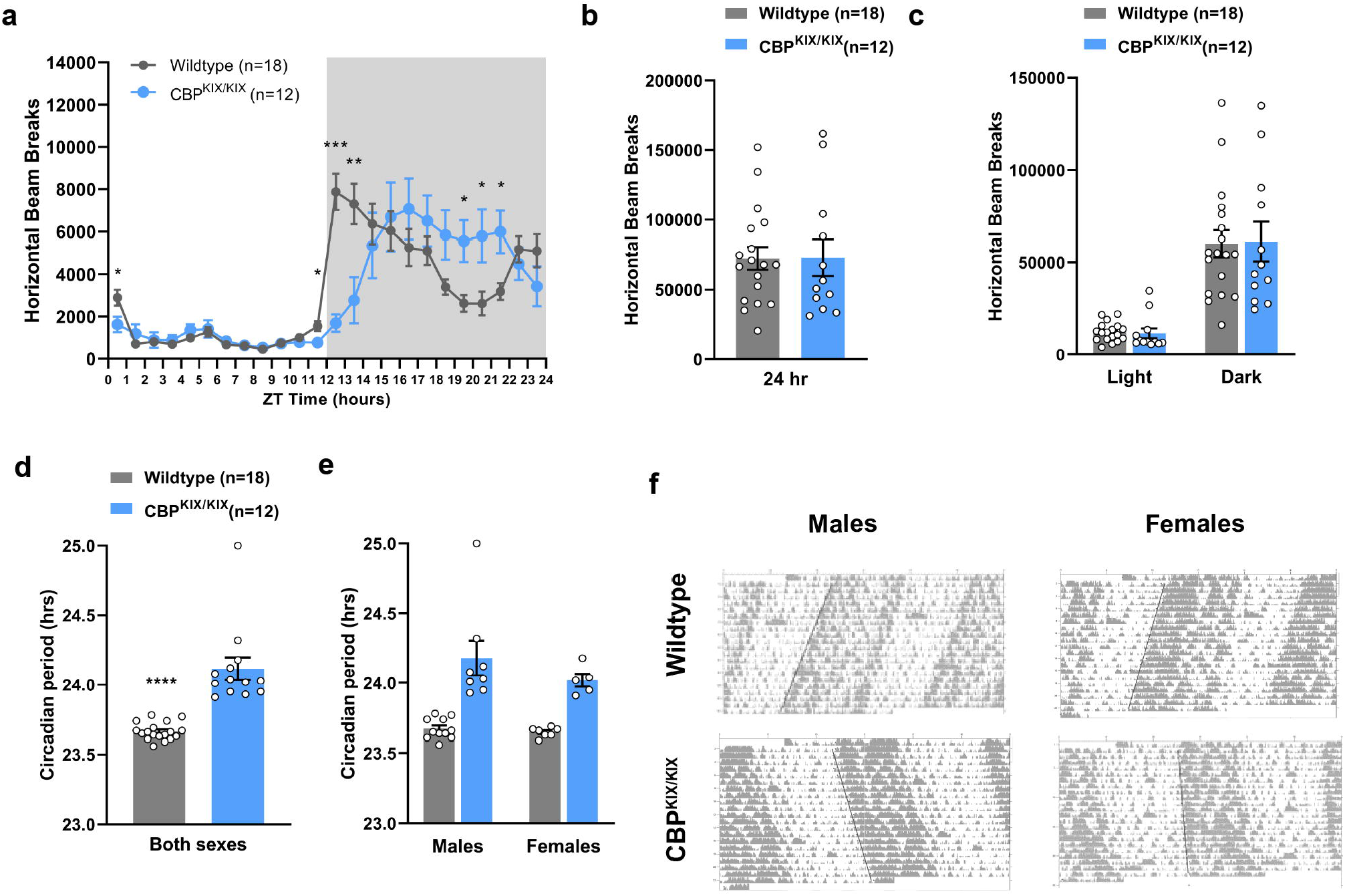
Male and female CBP^KIX/KIX^ mice have increased free-running periods compared to wildtype littermates. **(a)** CBP^KIX/KIX^ mice have delayed activity onset and peak activity in 12h:12h LD compared to wildtype littermates. Each data point represents 1 hour of binned activity data. Mixed Design ANOVA: Significant time x genotype interaction between CBP^KIX/KIX^ and WT mice (F_(23,644)_ = 8.26, p = 0.00003). **(b)** There are no significant differences in total activity levels between wildtype and CBP^KIX/KIX^ mice [t(28)=0.03183, p=0.9748]. **(c)** No significant differences were observed with the light and dark phase activity analyzed independently [Two-way ANOVA: no significant Genotype x Light/Dark interaction, F_(1, 56)_ = 0.02275, p =0.88]. **(d)** CBP^KIX/KIX^ male and female mice display longer free-running circadian periods in constant darkness compared to wildtype littermates (t _(29)_=6.550, p<0.0001). **(e)** Sex had no effect on circadian period of WT or CBP^KIX/KIX^ mice [Two-way ANOVA: Sex x Genotype interaction (F (1, 27) = 0.9139, p=0.3476), Main effect of genotype, F (1, 27) = 38.24, p<0.0001. **(f)** Representative actograms for males (left) and females (right).

To further probe the role of the KIX domain of CBP in the circadian clock, we investigated phase-shifting in CBP^KIX/KIX^ mice in response to light pulses in constant darkness. After 4 weeks in continuous darkness, one cohort of CBP^KIX/KIX^ mice and WT controls was given a 15-minute, 250 lux (80 lux at cage floor) phase delaying light pulse at CT14 for maximal phase delay (75). Circadian period length was recalculated and the phase shift in hours was calculated for each animal. Mice were allowed to again free-run in constant darkness for 8 days before receiving another light pulse, this time phase-advancing, at CT22 **(Fig 6a)**. There were no differences in response to the delaying (CT14) or advancing (CT22) phase shifts between CBP^KIX/KIX^ mice and WT controls (**Fig 6b-6d**). These data suggest that CBP-mediated transcription through the KIX domain is not necessary for circadian clock resetting, but rather is a modulator that plays a role in ensuring accurate timing of circadian transcriptional oscillations.

**Figure 6.**
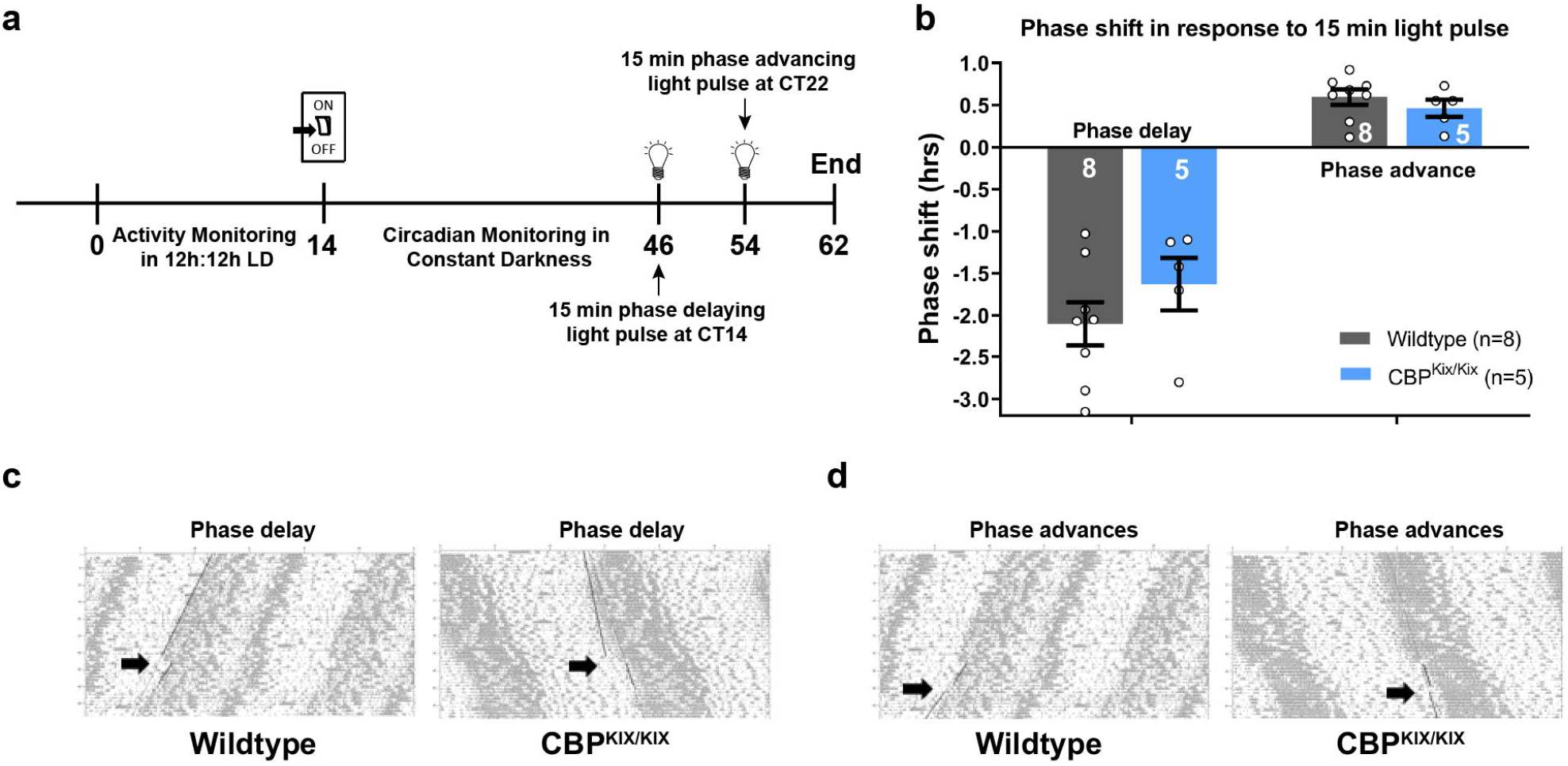
CBP^KIX/KIX^ mice exhibit normal responses to phase advances and phase delays. **(a)** Schematic representation of experimental timeline in days for circadian monitoring and light pulse experiments. **(b)** CBP^KIX/KIX^ and WT mice display comparable phase shifts when given either a phase delaying (CT14) or phase advancing (CT22) light pulse (Student’s t-test, phase delaying: (t_(11)_ = 1.15, *p* = 0.27; phase advancing: (t_(11)_ = 0.93, *p* = 0.37. **(c)** Representative actograms from CT14 phase delaying light pulse for wildtype (left) and CBP^KIX/KIX^ (right) mice. **(d)** Representative actograms from CT22 phase advancing light pulse for wildtype (left) and CBP^KIX/KIX^ (right) mice. *p<0.05, **p<0.01. ***p<0.001

## Discussion

CBP is a complex co-activator of transcription interacting with numerous transcription factors as well as functioning as a histone acetyltransferase important in long-term memory. To separate the multiple functions of CBP and provide clearer insight into the downstream targets of CREB-CBP transcriptional co-activation in memory, we used CBP^KIX/KIX^ mice with three point mutations in the KIX domain to analyze spatial memory following hippocampus-dependent learning and to identify downstream target genes. We found that CBP^KIX/KIX^ mice learned a complex spatial task, the MWM, with acquisition rates and repeat training induced performance improvements similar to WT mice. However, long-term spatial memory tested 24 hours following the last day of MWM training was not formed in CBP^KIX/KIX^ mice. These results are consistent with previously described *Cbp* mutant models with deficiencies in hippocampus dependent memory including contextual fear conditioning and object recognition memory (19–21). However, previous research using an adult specific forebrain *Cbp* knockout model found no deficits in long-term memory for MWM or contextual fear conditioning in the mutant mice (21). Potentially, compensation by other transcription factors occurs to make-up for the complete lack of forebrain CBP in these mutants, as these mice do not show deficits in numerous behavior tests including tests assessing anxiety, exploration, locomotor activity, motor coordination and learning (21). Thus, our results suggest a crucial role for the interaction of the KIX domain of CBP with transcription factors during the consolidation of long-term spatial memory.

Despite the lack of long-term spatial memory evident in CBP^KIX/KIX^ mice, these mice demonstrated memory recall in the one-hour probe test comparable to WT mice, and consistent with most previous research investigating short term-memory for contextual fear conditioning and object recognition in *Cbp* mutants (19, 20). It should be noted that deficits in associative short-term memory were found for contextual fear conditioning and novel object recognition in the forebrain *Cbp* knockout model that results in virtually complete depletion of CBP protein in excitatory neurons of the hippocampus and the cortex (76). In contrast, in the CBP^KIX/KIX^ mutant mice used in our studies, overall CBP protein levels and protein stability are not affected (19, 41). In CBP^KIX/KIX^ mice, CBP retains the potential to interact with non-KIX binding transcription factors to induce gene expression and CBP can also still function as an acetyltransferase. Although, CBP^KIX/KIX^ mice express the mutant form of CBP from developmental stages onward, we found only modest changes in baseline gene expression between CBP^KIX/KIX^ mice and WT littermates under homecage conditions.

To provide insight into the downstream transcriptional targets common to multiple hippocampus-dependent learning paradigms initiated through co-activation of CBP via the KIX domain, we performed deep RNA sequencing from the dorsal hippocampus of CBP^KIX/KIX^ mice and their WT littermates after training for either MWM or contextual fear conditioning. Consistent with the specificity of the mutations in the CBP^KIX/KIX^ mice, we found that CREB was the top-predicted upstream regulator of the differentially regulated genes in learning for CBP^KIX/KIX^ vs WT mice. To further support the hypothesis that compromised CREB-CBP interactions underlie the long-term deficits in hippocampal memory in CBP^KIX/KIX^ mice, we found that pCREB was decreased in CBP^KIX/KIX^ mice following three days of MWM training compared to WT mice. As Ser-133 phosphorylation is necessary for the binding of CREB to CBP and induces folding of the CREB kinase inducible domain (KID), the decreased binding of CREB to CBP may increase CREB vulnerability to phosphatase activity in the CBP^KIX/KIX^ mice as total CREB protein levels were not affected. Prior to binding CBP, the KID of CREB remains chiefly unstructured and is considered disordered with CBP binding increasing the helical structure of the KID region (77).

Our RNA-Seq results from the WT group comparison (learning versus homecage) supports the present literature that learning induces CREB and CREM target genes that both interact with CBP for its transactivation (12, 78, 79). Among the CREB downstream genes, we found several activity-dependent genes (such as *Arc*, *Egr1*, *Fos*, *Nr4a1*, *JunB*) that previously have been shown upregulated following hippocampus-dependent learning (31, 80, 81). It is noteworthy that some of these activity-dependent genes were downregulated in CBP^KIX/KIX^ mice, including *Nr4a1*. The role of the Nr4A sub-family has been well characterized in memory. Reducing Nr4A function in the hippocampus impairs long-term memory (82–84), while the activation of Nr4A family transcription factors enhances memory in young (85) and aged mice (86, 87). Pharmacological activation of CBP in a mouse model of Alzheimer’s disease rescues *Nr4A* gene expression and long-term memory (31).

Notably, we found that genes involved in circadian rhythms were significantly downregulated in the dorsal hippocampus of CBP^KIX/KIX^ mice following learning including *Per1, Per2, Per3, Dbp, Nr1d1, Nr1d2,* and *Ciart/Chrono*. IPA analysis revealed the major upstream regulators of the downregulated circadian genes in CBP^KIX/KIX^ mice following learning were CREB, CLOCK and BMAL1 emphasizing the interaction between CBP and the circadian oscillator. Although the suprachiasmatic nucleus (SCN) houses the master circadian clock in mammals, circadian rhythms and circadian genes in the hippocampus have been associated with memory. Performances in hippocampus-dependent tasks including the radial arm maze, spontaneous alteration, novel object location, contextual fear conditioning, passive avoidance and MWM are time of day dependent (reviewed in (88)). The formation and persistence of long-term hippocampal memory has been associated with circadian rhythms in cAMP and MAPK signaling through CREB-mediated transcription (89). Hippocampal PER1 has been postulated to function as a gatekeeper conveying time of day information to signaling pathways involved in memory formation through the regulation of CREB phosphorylation (57, 59). While these previous studies indicate that the circadian clock regulates CREB mediated transcription in the hippocampus to affect memory formation, our data suggests reciprocal interactions with CBP transcriptional co-activation mediating *Per* gene expression after learning. Our research suggests that in the hippocampus, *Per1* and *Per2* transcription is dependent upon CBP transcriptional co-activation similar to the CBP-CREB-dependent regulation of *Per* genes seen in the SCN (90, 91).

Previously, *in vitro* research has indicated that CBP modulates circadian rhythms and interacts with BMAL1 in phase resetting of the circadian clock (40). Moreover, siRNA knockdown of CBP significantly diminishes circadian oscillations in cultured cells (40). However, no previous studies in mammalian models have examined the effects of CBP mutations on circadian rhythms *in vivo.* We found that under diurnal conditions, CBP^KIX/KIX^ mice exhibited delayed activity onset with peak activity occurring significantly later in the night compared to sex-matched controls. In constant darkness, both male and female CBP^KIX/KIX^ mice have significantly longer free-running periods. During preparation of this manuscript, high resolution structural research identified direct interactions between the C-terminal transactivation domain of BMAL1 with the KIX domain of CBP (92). Thus, it appears possible that CBP affects circadian clock function through independent interactions with circadian transcription factors, particularly BMAL1, in addition to transcriptional co-activation with phospho-CREB (Ser133). The role of CBP in the circadian clock appears highly conserved. In *Drosophila*, CBP has been shown important in circadian gene transcription regulating the CLOCK/CYCLE activation of transcription (93–96). Similar to what we observed in mice, when CBP levels are decreased in circadian pacemaker neurons, the period is lengthened in *Drosophila* (95). Moreover, the regulation of CBP levels appears to be a critical factor in the maintenance of normal circadian function as overexpression of CBP induces behavioral arrhythmicity in *Drosophila* (93, 96).

Despite the temporally delayed activity pattern and longer free-running rhythms seen in CBP^KIX/KIX^ mice, we found no difference between phase resetting in mutant mice and WT by light pulses delivered in the early or late night. The dissociation between the impact of KIX domain mutations on different functions of the circadian clock is consistent with the multiple domain nature of the CBP protein. For phase resetting of the circadian clock, the interactions between CBP and BMAL1 at the *Per1* promoter are thought to be primarily reliant upon calcium-dependent PKC signaling following resetting stimuli and not CREB (40). Moreover, CBP has been postulated to interact with BMAL 1 through the PXDLS motif (Pro-X-Asp-Leu-Ser), a conserved motif in multiple circadian proteins and present in CBP (64). Our *in vivo* studies differentiating the impact of KIX mutations on period length and phase resetting support the hypothesis that different signaling pathways trigger CBP involvement in these two functions. Since phase shifts in response to light have been shown to be, at least in part, CREB-dependent, CREB also may act through binding to other co-activators such as CREB-regulated transcription coactivator (CRTC1) in phase resetting via Per1 in the SCN (97, 98).

Our research on the CBP KIX domain provides significant insight into two processes that significantly impact healthy aging, memory and circadian rhythms. Although overall CBP protein levels appear unchanged with aging in one rat model (99), potentially, changes in CREB-CBP interactions through the KIX domain could be linked to age-related memory impairments. Mutations in CBP have been associated with neurodegenerative diseases including Huntington’s Disease (32, 33) and Alzheimer’s disease (34–36). Post-mortem brain analysis in humans and animal models have found decreased CREB expression associated with many neurological and neurodegenerative disorders including Alzheimer’s disease, Parkinson’s disease, bipolar disorder and schizophrenia (31, 100–102). In a mouse model of Alzheimer’s disease, viral delivery of CBP restores pCREB levels and mitigates cognitive impairments (103). Our research highlights the role the CBP KIX domain plays in the regulation of circadian gene expression and circadian activity as well as its role in long-term memory. Maintaining robust circadian rhythms is also essential for healthy aging. Age-related changes in circadian function include the dampening of molecular circadian rhythms, changes in free-running period, and impairments in the synchronization and coordination of circadian oscillators across tissues and organs (reviewed in (104)). The longer free running period in CBP^KIX/KIX^ mice with delayed activity onset is reminiscent of the circadian phenotypes seen in aging mice. Aged mice have significantly longer free-running periods and exhibit a delay in activity onset after dark compared to younger adult mice (105). Recent evidence suggests that circadian rhythm disruptions may contribute to increased risk or aggravate disease pathology for neurodegenerative diseases including Alzheimer’s disease and Parkinson’s disease (106–111). Additionally, recent research suggests that hippocampal Per1 specifically may be critical to age-related cognitive impairments (58). The CBP KIX domain appears to represent a potential target for understanding the effects of circadian and memory impairments that occur with aging. With life expectancy predicted to continue rising (112), identifying the regulatory mechanisms and molecular processes underlying the formation of memory is a necessary step for the development of future therapies and treatments for memory impairments.

## Materials and methods

### Animals

The CBP^KIX/KIX^ mice for experiments were generated from heterozygous mating (from C57BL/6J genetic background), with wildtype (WT) littermates used as controls. Mice were approximately 3 months old at the start of experiments. Male and female mice were used in experimental and control groups. Mice were housed in groups of 2-3 under 12-h light/12 h-dark cycle in a temperature- and humidity-controlled room (22°C and 55 +/− 5%, respectively) with *ad libitum* access to food and water. All behavioral training and testing (with the exception of circadian monitoring) was performed during the light portion of the cycle (lights on at 7:00 A.M.) at the same time of day. Experimental protocols and animal care at the University of Strasbourg were in compliance with the institutional guidelines laws and policies (the European Parliament 2010/63/UE of September 22, 2010). All experiments at the University of Pennsylvania were approved by the University of Pennsylvania Institutional Care and Use Committee (IACUC protocol 804407) and conducted in accordance to National Institute of Health guidelines.

### Morris water maze

Evaluation of spatial memory was performed using the Morris Water Maze task (MWM) as described in Chatterjee et al., 2013 (30). For the spatial memory tests, two independent groups of mice CBP^KIX/KIX^ and their WT littermates were trained for five days. Each training day consisted of four trials of maximum duration of 60 sec in each trial to locate a hidden platform under the surface of water using the visual cues present in the room. After the last training session, the platform was removed, and each group of animals was tested (probe test) to measure short-term memory 1 h later or 24 h later to measure long-term memory. During the probe test, the mice were placed in the opposite quadrant with respect to the target quadrant and allowed to swim for 60 s. Spatial memory during the probe test was quantified by measuring the amount of time spent by the mice searching in the target quadrant versus the average time spent in the three other quadrants. For the biochemical studies, mice were trained for three consecutive days. Three days of training using this protocol has previously been shown sufficient to induce CBP-dependent gene expression changes in the dorsal hippocampus (113). One hour after the last trial on the 3^rd^ day, the dorsal hippocampus was immediately dissected, frozen in liquid nitrogen and stored at −80°C.

### Contextual fear conditioning

Contextual fear conditioning was performed as previously described (114) with mice handled one minute per day for three consecutive days prior to fear conditioning training. On the training day, the mice received a single 2 s, 1.5 mA foot-shock culminating 2.5 min after placement into the chamber. Mice were removed from the chamber 30 s after receiving the foot shock and returned to their homecages. A separate group of mice serving as homecage controls were handled but did not receive the electric shock.

### Microarray

Microarray analysis was performed with Affymetrix MOE 430v2 arrays probed with labelled RNA from the two hippocampi of each animal as a single biological replicate. RNA was obtained using a TRIzol-chloroform extraction followed by RNAeasy (Qiagen) isolation. RNA quality was assessed with the Agilent Bioanalyzer. RNA amplification and labelling were performed with the NuGen ovation system. After hybridization and array reading at the Penn Microarray Facility, intensity scores were calculated by the Robust Multiarray Average (RMA) method (115), and quality control was performed using array Quality Metrics (116) to ensure that differences in signal intensity did not bias analysis. Following the RMA, patterns from Gene Expression (PaGE v5.1) analysis was implemented in Perl to assess confidence in changes in probe set expression level between CBP^KIX/KIX^ and wildtype littermates. This algorithm uses permutations of the input data to empirically determine the False Discovery Rate (FDR) for a given T-statistic level, allowing assignment of FDR values to individual probe sets based on a modified T-statistic. FDR values were computed with 200 permutations, and multiple instances of FDR calculation were run to assess confidence levels (1-FDR) from 80% to 95%.

### RNA extraction for RNA-seq library preparation and sequencing

Dorsal hippocampus was quickly dissected, placed into RNA later (Qiagen Valencia, CA) and frozen in liquid nitrogen. Tissues were homogenized in QIAzol Lysis Reagent (Invitrogen) and mixed in chloroform. RNA in aqueous-phase was separated in phase-lock tubes following centrifugation at 14,000 g for 15 mins. RNA was extracted using RNAeasy kit (Qiagen) according to the manufacturer’s instruction. Samples were DNase treated using the RNase-Free DNase kit (Qiagen) off-column. Samples were ethanol precipitated and resuspended in RNAse-free water. Samples with an OD 260/280 and OD 260/230 ratio close to 2.0 and RNA integrity number (RIN) above 8 were selected for library preparation. RNA library from homecage and fear conditioned mice were prepared at the PGFI sequencing core at the University of Pennsylvania using the TruSeq sample preparation kit (Illumina San Diego, CA) according to the manufacturer’s instructions with polyA selection. Libraries were size selected and quantified by qPCR (KAPA Biosystems Boston, MA), and sequenced. Barcoded libraries were sequenced in Illumina HiSeq 2000 in paired-end 100 bp reads. Three libraries were sequenced per lane on an Illumina HiSeq 2000 at the sequencing core at the University of Pennsylvania.

RNA libraries from homecage and MWM mice were prepared at the Iowa Institute of Human Genetics (IIHG), Genomics Division, using the Illumina TruSeq Stranded Total RNA with Ribo-Zero gold sample preparation kit (Illumina, Inc., San Diego, CA). Library concentrations were measured with KAPA Illumina Library Quantification Kit (KAPA Biosystems, Wilmington, MA). Pooled libraries were sequenced on Illumina HiSeq 4000 sequencers with 150-bp Paired-End chemistry (Illumina) at the IIHG core.

The RNA seq data have been deposited in NCBI’s Gene Expression Omnibus and are accessible through GEO Series accession number GSE151681 (https://www.ncbi.nlm.nih.gov/geo/query/acc.cgi?acc=GSE151681). The code for analyses and figures related to RNA-seq data can be accessed through GitHub (github.com/ethanbahl/chatterjee2020_cbpkix).

#### RNA-seq analysis

Sequencing data was processed with the bcbio-nextgen pipeline (https://github.com/bcbio/bcbio-nextgen). The pipeline uses STAR (117) to align reads to the genome and quantifies expression at the gene level with featureCounts (118). All further analyses were performed using R (119). For gene level count data, the R package EDASeq was used to adjust for GC content effects (full quantile normalization) and account for sequencing depth (upper quartile normalization) (120) **(Supplemental Fig S4 and S5)**. Latent sources of variation in expression levels were assessed and accounted for using RUVSeq (RUVr) (121)(**Supplemental Fig S6 and S7)**. Appropriate choice of the RUVSeq parameter *k* was determined through inspection of P-value distributions, mean-difference plots, RLE plots, and PCA plots. Specifically, values of *k* were chosen where P-value distributions showed an expected peak below 0.05 (122) and experimental groups were separable with three principal components. Differential expression analysis was conducted using edgeR (123, 124).

We performed two differential expression analyses to identify genes responsive to learning in wildtype mice, and genes responsive to the CBP^KIX/KIX^ genotype following learning. Based on criteria listed above, we used *k* values of 3 and 4 for the two analyses. Data generated from two separate learning paradigms were included in the differential expression models. Effects specific to a particular experiment (i.e., MWM or contextual fear conditioning) were accounted for in the models, thus revealing genes reproducibly responsive to learning or the CBP^KIX/KIX^ genotype across multiple paradigms of learning.

To externally validate our analysis, we used previously published research on gene regulation in the hippocampus following chemically-induced neuronal activation (55). We created a positive control set (**Supplemental Table 4 with list of positive controls**) using genes in the top decile of adjusted P-values (adjusted P < 0.05). We performed a Fisher’s exact test on a two-dimensional contingency table, with rows indicating whether a gene was responsive to learning in our study, and columns indicating whether the gene was in the positive control set. Using this approach, we also tested the set of genes responsive to the CBP^KIX/KIX^ genotype, after learning, for enrichment of genes responsive to learning in WT animals.

#### Upstream regulator and pathway analysis

Upstream regulator analysis was performed using the QIAGEN’s Ingenuity® Pathway Analysis (Qiagen Redwood City, CA, USA, www.qiagen.com/ingenuity) as described in Lorsch et al (125). Global enrichment network analysis from KEGG pathway database were performed using the free online NetworkAnalyst software 3.0 (https://www.networkanalyst.ca).

### Activity Monitoring

Activity monitoring was performed as previously described in a separate group of mice from the RNA-seq / microarray/ behavioral experiments (65, 126). Briefly, mice were individually housed inside light- and noise-attenuating chambers (22” × 16” × 19”, Med Associates, St. Albans, VT) equipped with a 250-lux light source (80 lux at cage floor) and fan for ventilation. Each cage was placed within a system of infrared beams spaced 0.5 inches apart that provided two horizontal infrared grids at 0.75” and 2.75” above the cage floor. Mice were allowed to acclimate to the activity monitoring chambers for one week before the start of activity monitoring. Infrared beam break counts were acquired at 10 second intervals for 7 days on a 12 h:12 h LD schedule to measure both horizontal and vertical (rearing) activity across the entire diurnal cycle. Activity counts were binned into 1-hour intervals and averaged over the 7 days. After the final day of activity monitoring in LD cycles, lights were switched off and animals were allowed to free-run in 24 h constant darkness (DD) for 4 weeks with activity counts compiled every minute. Circadian period (tau) was calculated from Day 2 until the end of DD using ClockLab software (Actimetrics).

### Light Pulses

Following 4 weeks of constant darkness, one cohort of mice (n=8 WT and n=5 CBP^KIX/KIX^) received a 15 min phase delaying 250 lux (80 lux at cage floor) light pulse at CT 14. Mice were allowed to recover for 8 days in 24 h constant darkness and the phase shift was measured. After the 8 days in continuous darkness, a second 15 min phase advancing light pulse was given at CT 22. The phase angles of activity delay and advancement were calculated using ClockLab software (Actimetrics).

### Statistical analyses

All statistics were performed using Graphpad Prism (V8.1), except for circadian analysis which was performed using SPSS for Windows (v 24.0) and the bioinformatics analysis. For MWM, performance recorded during acquisition (latency to the platform) was evaluated using a one-or two-way ANOVA with repeated measures when appropriate. Escape latencies during acquisition were analyzed using two-way ANOVA, followed by Sidak’s multiple comparisons test. Probe trial performance was also analyzed using two-way ANOVA, followed by Sidak’s multiple comparisons test comparing the time spent in the target quadrant (T) and the time spent in the other three quadrants (Q1-3). Homecage activity was analyzed using a Mixed ANOVA with genotypes (WT and CBP^KIX/KIX^) as the between-subjects factor and time as the within-subject factor. Post hoc multiple comparisons were performed. Multivariate ANOVAs (MANOVA) were used to analyze the activity counts in the light and dark phases, with alpha corrected for multiple ANOVAs and set at α = 0.05/2. For total activity over the 24 hour day and light pulse analysis, Student’s t-test was applied. Results were expressed as means +/− SEM. Values of p < 0.05 were considered as statistically significant.

## Acknowledgements

This work was supported by the National Institute of Health R01 MH087463 and R01 MH 087463-08S1 to T. A., Centre National de la Recherche Scientifique, the University of Strasbourg and the Indo-French Centre for the Promotion of Advanced Research (Grant #4803-3 to A.-L. B). T. A. was also supported by the Roy J. Carver Chair in Neuroscience at the University of Iowa.

## Footnotes

CCA’s present address is: Department of Psychiatry and Behavioral Sciences, Stanford University, Stanford, California, USA.

SGP’s present address is: DNA Electronics, San Diego, California, USA.

JDH’s present address is: Department of Cellular Neuroscience, Neurodegeneration, and Repair, Yale University, New Haven, Connecticut, USA.

