## Supplemental information for "The CBP KIX domain regulates long-term memory and circadian activity"

### Supplemental Figures and legends

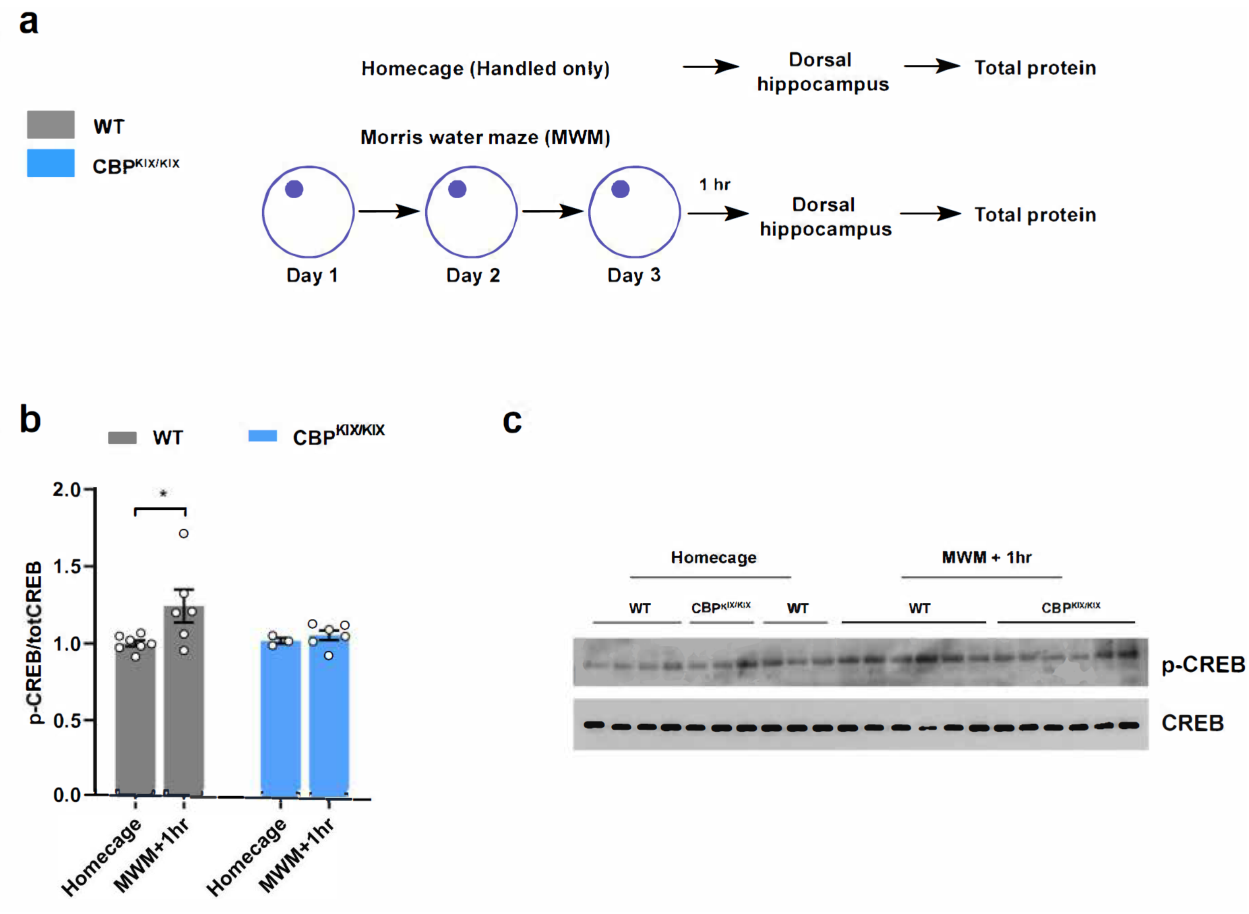


**Figure S1. Learning induced CREB phosphorylation at S133 is decreased in CBP$^{KIX/KIX}$ mice. (a)** Scheme of the experiment**. (b)** CREB phosphorylation at S133 is significantly increased after MWM training ([Unpaired t-test: $t_{(11)}$=2.407, p=0.0348, WT HC Vs WT MWM], while no such enhancement were observed in CBP$^{KIX/KIX}$ mice [Unpaired t-test: $t_{(7)}$=0.7450, p=0.4805]. **(c)** Western blot showing CREB and p-CREB expression.

**Supplemental Fig S2**

| Gene | Probe set | HC confidence | HC KIX/WT ratio |
|---|---|---|---|
| Crebbp | 1459804_at | 99.5% | 0.63 |
| Crebbp | 1436983_at | 0.0% | 1.06 |
| Crebbp | 1435224_at | 0.0% | 1.01 |
| Crebbp | 1434633_at | 0.0% | 1.02 |

**Figure S2. CBP gene expression is unaltered in CBP$^{KIX/KIX}$ mice.** The four probe sets for CBP, official gene symbol Crebbp, are shown. Only the probe set that hybridizes to the KIX domain (1459804_at) shows evidence of reduced signal intensity in CBP$^{KIX/KIX}$ mice. The approximately 40% reduction in signal only for this probe set is likely due to the three base pair mutations introduced by the KIX domain mutation.

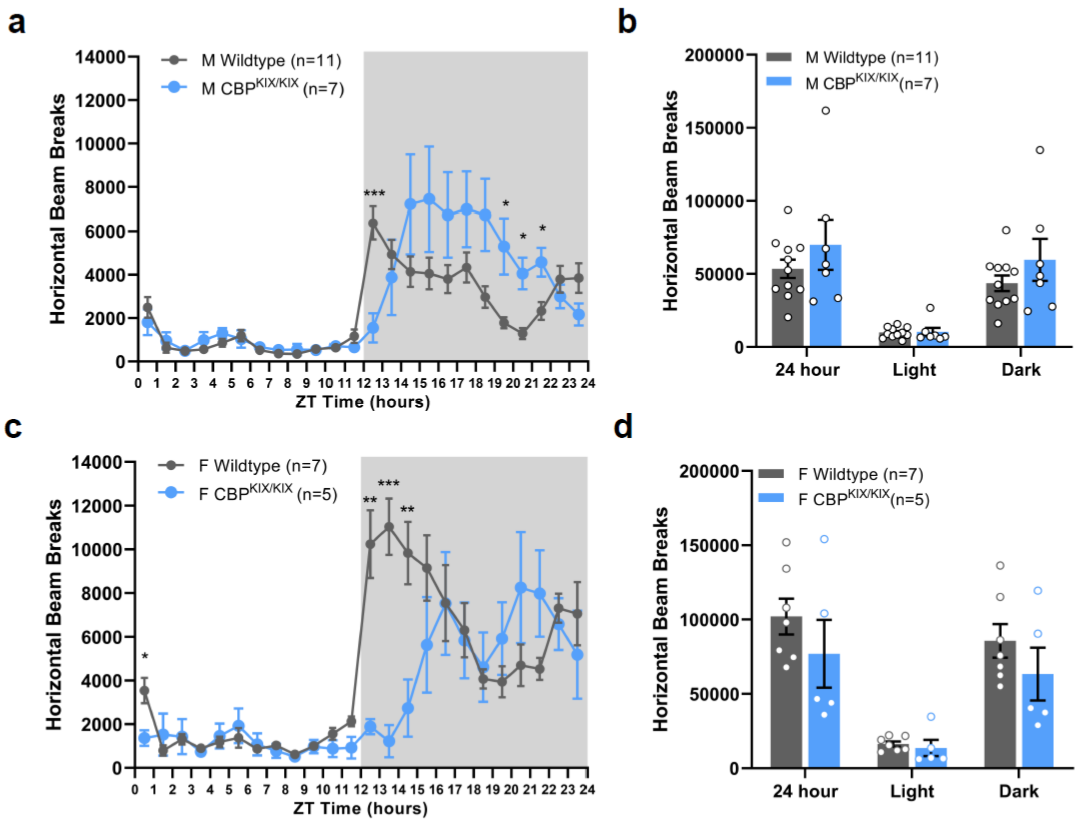


**Figure S3. Delayed peak diurnal activity in male and female CBP$^{KIX/KIX}$ mice. A,B.** CBP$^{KIX/KIX}$ males have delayed activity onset and peak activity in 12h:12h LD compared to wildtype littermates **(a)** (time*genotype, F(23,368) = 6.13, p = 0.002), without exhibiting differences in total ambulation **(b)** (t(16) = 1.16, p= 0.307). **c, d.** CBP$^{KIX/KIX}$ females also have delayed diurnal activity patterns compared to wildtype (C) (F(2,23) = 7.95, p < 0.001), while total movement is unaltered between CBP$^{KIX/KIX}$ females and wildtype controls (d) (t(10) = 1.11, p = 0.316). *p<0.05, **p<0.01. ***p<0.001

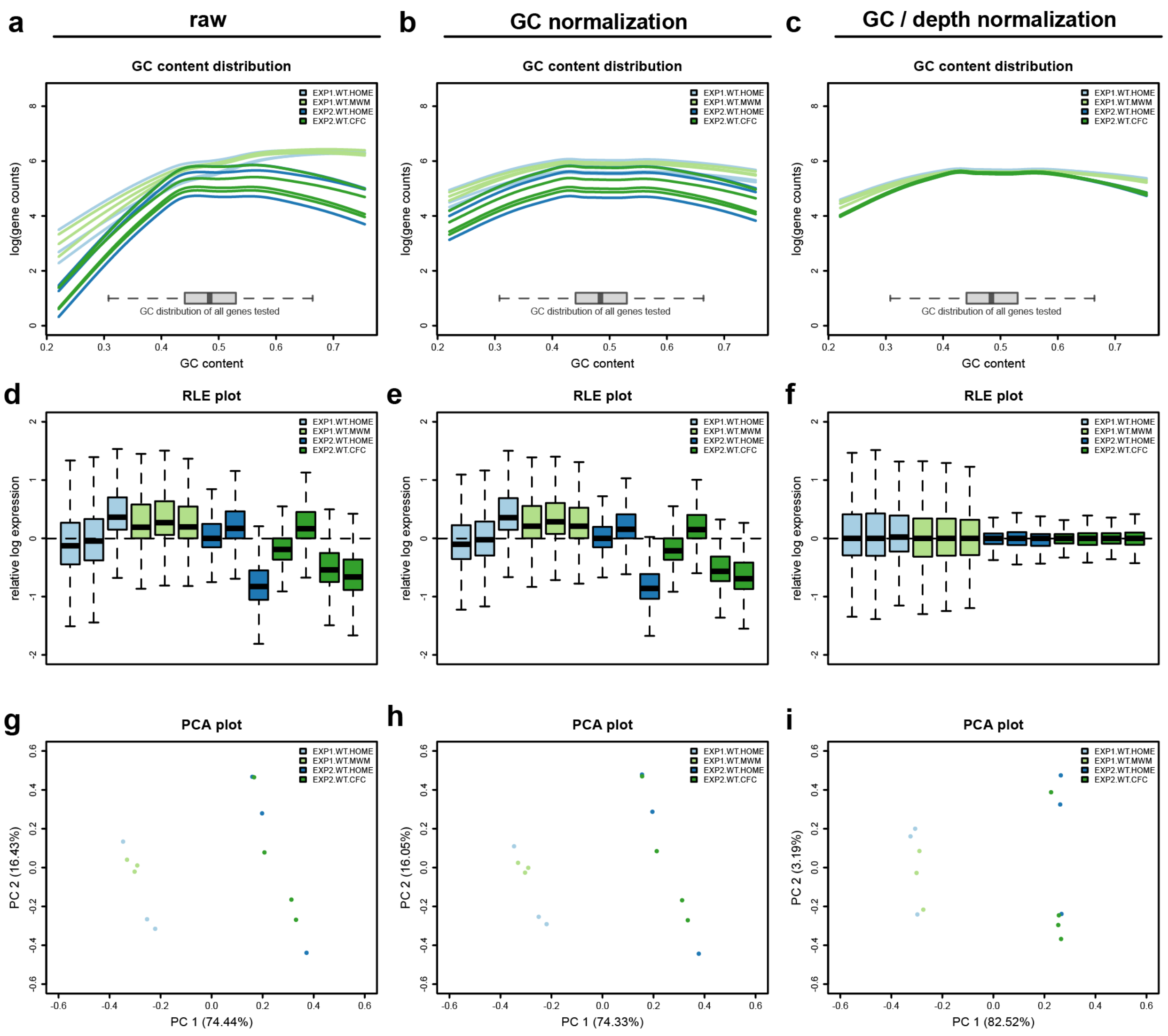


**Supplemental Figure S4**

**Figure S4. Normalization of differences in GC content distribution and sequencing depth using EDASeq for comparisons of RNAseq studies after learning in wildtype mice and CBP$^{KIX/KIX}$ mice.** Distributional differences in GC content and variability in sequencing depth are sources of technical variability in RNA-sequencing data. **a-c**, GC content distributions before normalization (**a**), after full quantile GC content normalization (**B**), and upper quartile sequencing depth normalization (**c**). **d-f**, Relative log expression (RLE) plots before normalization (**d**), after full quantile GC content normalization (**e**), and upper quartile sequencing depth normalization (**f**). **g-i**, Principal component analysis (PCA) plots before normalization (**g**), after full quantile GC content normalization (**h**), and upper quartile sequencing depth normalization (**i**).

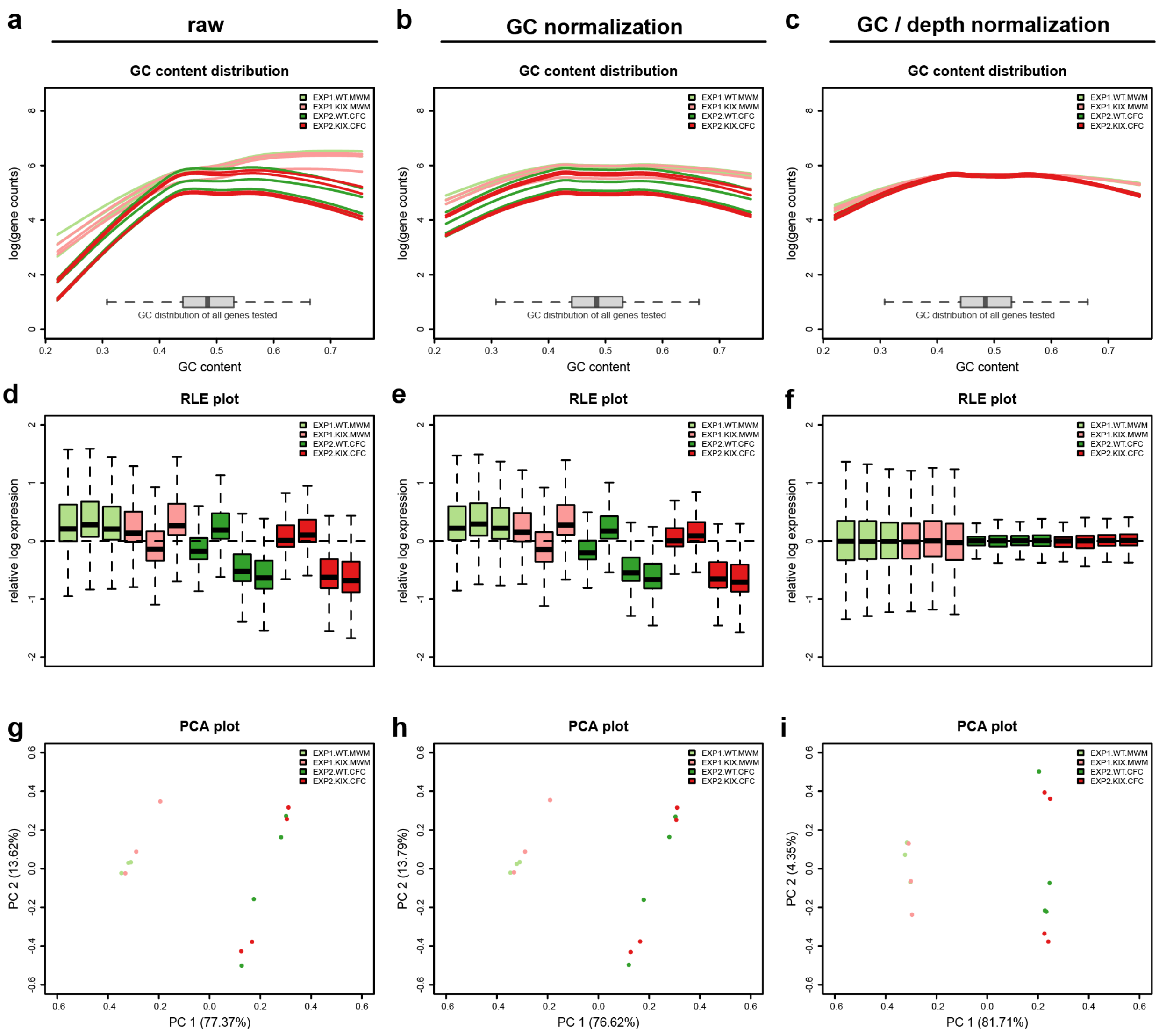


**Supplemental Figure S5**

**Figure S5. Normalization of differences in GC content distribution and sequencing depth using EDASeq for comparisons of RNAseq studies between homecage wildtype mice and trained wildtype mice. a-c**, GC content distributions before normalization (**a**), after full quantile GC content normalization (**b**), and upper quartile sequencing depth normalization (**c**). **d-f**, Relative log expression (RLE) plots before normalization (**d**), after full quantile GC content normalization (**e**), and upper quartile sequencing depth normalization (**f**). **g-i**, Principal component analysis (PCA) plots before normalization (**g**), after full quantile GC content normalization (**h**), and upper quartile sequencing depth normalization (**i**).

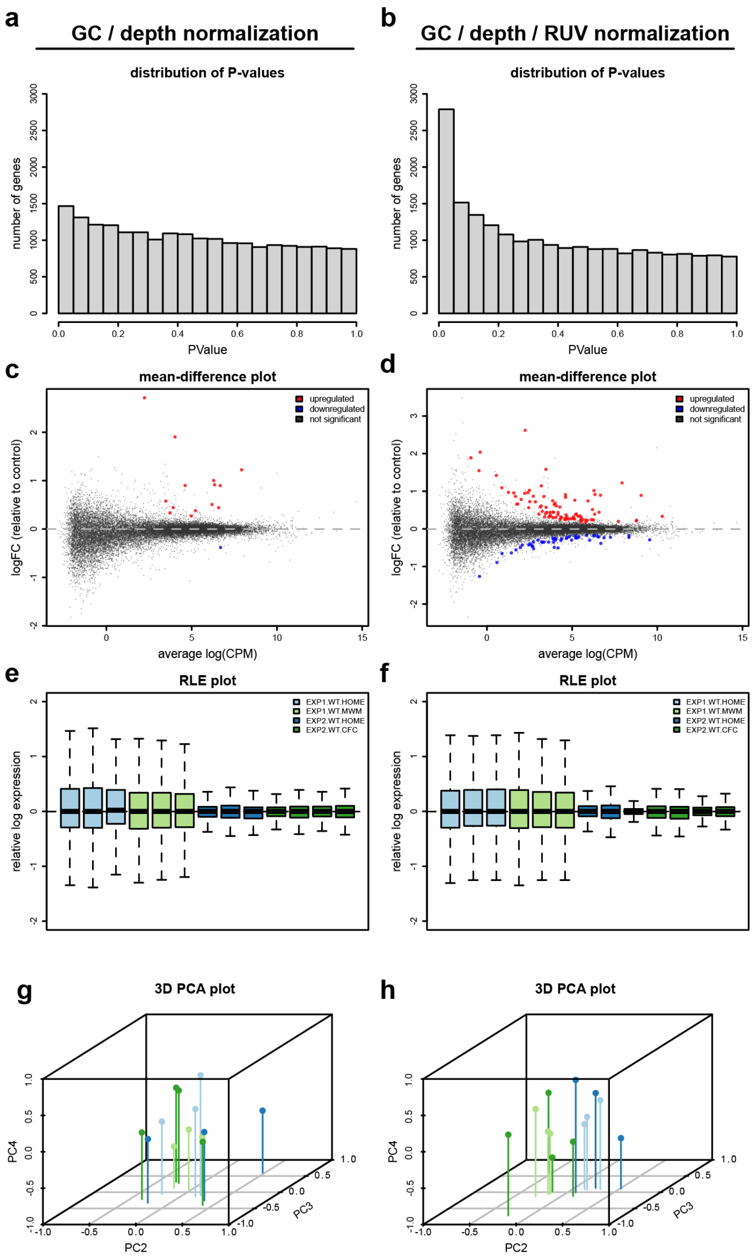


**Figure S6. RUV normalization for analysis of RNAseq after learning in wildtype mice and CBP$^{KIX/KIX}$ mice.** RUV normalization removes unwanted variation that dwarfs biological signal in RNA-sequencing data. **Left.** Without applying RUV normalization, uncorrected P-values from the differential expression analysis are uniformly distributed and few genes are found to be statistically significant at false discovery rate (FDR)<=0.05 after multiple testing correction (**a,c**). RLE and PCA plots reveal traditional normalization approaches fail to allow separation of biologically meaningful groups using three principal components* (**e,g**). **Right.** Applying RUV normalization increases power to detect statistically significant differences in gene expression (**b,d**), and allows separation of experimental groups (blue-green, green-red) of interest using three principal components* (**f,h**). **For 3-dimensional PCA plots, the first component represents experimental batch (correlation coefficient > 0.99). As experimental batch is directly accounted for in the differential expression model, we excluded it here and considered PC 2-4 for visualization and the assessment of RUV normalization*

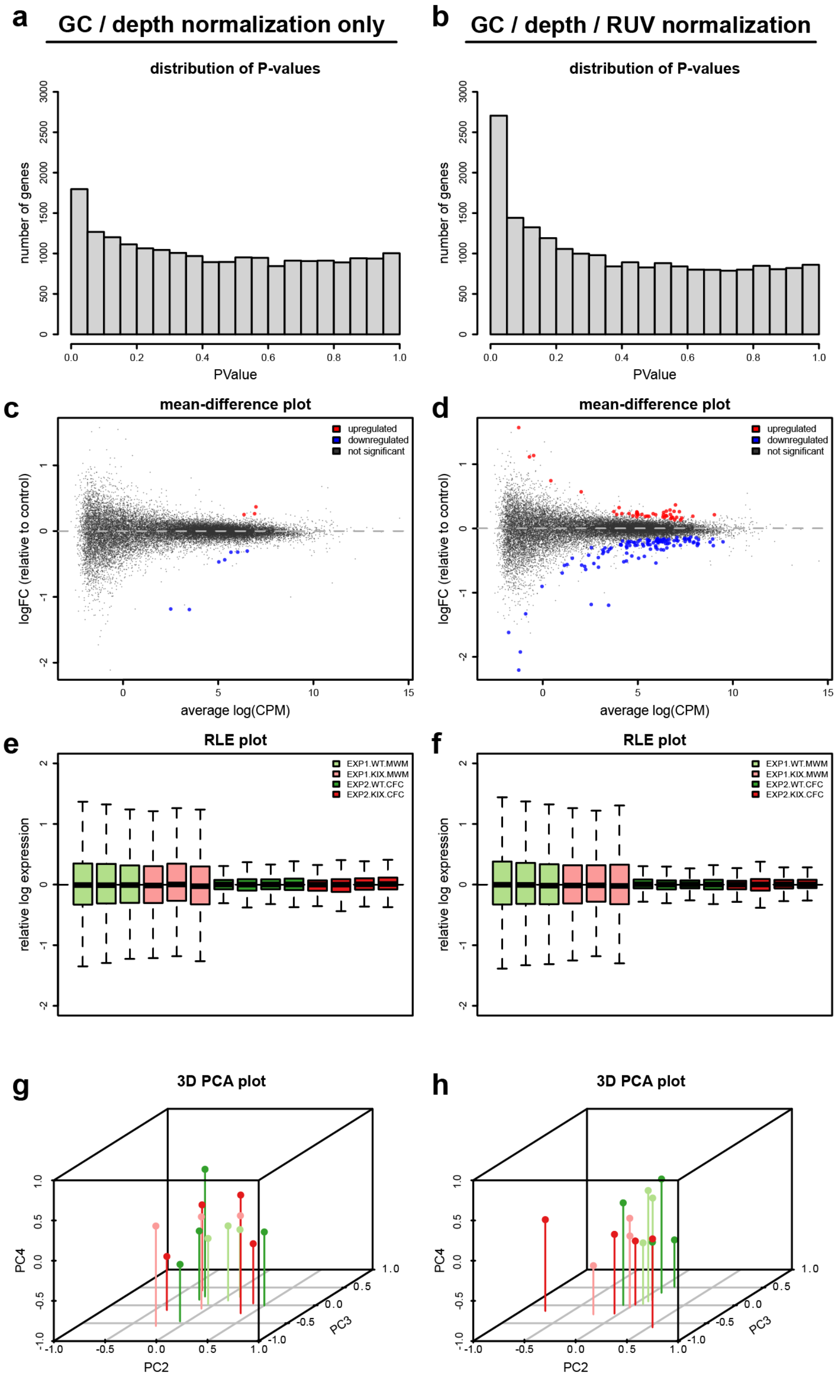


**Figure S7. RUV normalization for analysis of homecage and trained wildtype mice RNAseq experiments. Left.** Without applying RUV normalization, uncorrected P-values from the differential expression analysis are uniformly distributed and few genes are found to be statistically significant at false discovery rate (FDR)<=0.05 after multiple testing correction (**a,c**). RLE and PCA plots reveal traditional normalization approaches fail to allow separation of biologically meaningful groups using three principal components* (**e,g**). **Right.** Applying RUV normalization increases power to detect statistically significant differences in gene expression (**b,d**), and allows separation of experimental groups (blue-green, green-red) of interest using three principal components (**f,h**).

**Supplemental Tables**

**Supplemental Table 1.** List of Genes Differentially Expressed in $CBP^{KIX/KIX}$ mutants compared to wild-type littermates after spatial learning.

**Supplemental Table 2**. List of the upstream regulators of the DEGs in $CBP^{KIX/KIX}$ mutants

**Supplemental Table 3**. List of Genes Differentially Expressed in wild-type mice after spatial learning compared to homecage.

**Supplemental Table 4.** Positive controls for validation of differential gene expression analysis.

**Supplemental Table 5**. List of the upstream regulators of the DEGs in wild-type mice following learning.

**Supplemental Table 6**. High confidence (80%) changes in home-cage gene expression in $CBP^{KIX/KIX}$ mice.
